# Classic and novel features of the dentate gyrus vascular niche quantified through 3D vessel reconstruction

**DOI:** 10.64898/2026.10.20.760295

**Authors:** India Carter, Bryon M. Smith, Elizabeth D. Kirby

## Abstract

Adult neurogenesis, the continued generation of new neurons throughout life, supports a variety of emotional and cognitive functions in mammals. Lifelong neurogenesis occurs in the dentate gyrus (DG) of the hippocampus, where a neurogenic niche supports the survival and growth of neural stem cells (NSCs), which divide to generate new neurons. A key component of this niche is the surrounding vasculature, thought to give NSCs access to circulating support molecules, oxygen, and nutrient metabolites. The DG vasculature is strikingly variable across layers. Vascular density and linearity, for example, have been noted to differ substantially in the different cell layers that make up the DG. Most of these prior descriptions of the DG vasculature have been qualitative, and other important aspects of vessel morphology, like vessel diameter, have not been previously reported to our knowledge. To provide a more complete and quantitative description of the adult DG vasculature, we used a 3D imaging technique to fully characterize the vasculature of each DG cellular layer. Our 3D reconstructions of DG blood vessels provide a quantitative method for measuring several known, qualitative features of the neurogenic niche, such as blood vessel density and tortuosity. In some cases, we reproduce qualitative observations, such as affirming that vessel density is high in the subgranular zone (SGZ) where NSCs reside and low in the granule cell layer. In other cases, our findings go counter to what qualitative observations suggest. For example, the SGZ is often described as having uniquely linear blood vessels, but we found that linearity was similar between the SGZ and hilus layer. We also report new measures of features of DG layer vascular architecture: vessel diameter and branching. Together, these findings offer a global, quantitative characterization of the DG vascular niche in adult mice. Beyond providing more detailed characterization of DG vasculature, we also anticipate our methods and estimates of variance in mice will be useful for investigators wishing to plan their own quantitative studies of changes in the vascular structure of the DG.

## Introduction

Neurogenesis, the process of generating new neurons, persists into adulthood within select niches in most mammalian species (Ming and Song, 2011). In adult mice, there are two separate brain areas that undergo adult neurogenesis: the subventricular zone (SVZ) in the lateral ventricles and the subgranular zone (SGZ) in the dentate gyrus (DG) of the hippocampus (Alvarez-Buylla and Lim, 2004; Ming and Song, 2011). Within the SGZ, adult neurogenesis begins with radial glial like neural stem cells (RGL-NSCs), which can divide to generate proliferative progeny, known as intermediate progenitor cells (IPCs) (Olpe and Jessberger, 2023). Daughter IPCs then generate progeny that differentiate into neuroblasts and eventually into granule neurons. The new dentate granule neurons incorporate into the circuitry of the DG and become involved in learning and memory, as well as affect regulation (Aimone et al., 2014; Li et al., 2023; Alonso et al., 2024). Dysregulation of adult hippocampal neurogenesis has been suggested to be linked to several neurological diseases like Alzheimer’s disease and other hippocampal memory impairments (Toda et al., 2019).

The niches of the adult neurogenic regions are critical to supporting healthy neurogenesis that continues through adulthood. A particularly crucial component of the DG stem cell niche is the vasculature. Within the DG, the vasculature has been widely reported to be especially dense and planar (Palmer et al., 2000). RGL-NSC and IPC somas in the SGZ reside in close proximity to these densely arranged blood vessels, a relationship that emerges progressively over postnatal development and appears to be driven by stem and progenitor cell motility (Dause et al., 2024, 2025; Devasthali et al., 2025). NSCs also possess an apical process which extends through the DG to wrap around vessels found in the inner molecular layer (ML) (Moss et al., 2016; Licht et al., 2020). This unique arrangement of stem/progenitor cells and vasculature in the DG is hypothesized to support adult neurogenesis in several ways, such as by providing scaffolding for progenitors and neuroblasts to migrate tangentially through the DG, as well as by providing NSCs and IPCs access to oxygen and circulating support molecules such as growth factors and nutrient metabolites (Bovetti et al., 2007; Whitman and Greer, 2009; Licht et al., 2020).

Despite how critical the vascular networks are to preservation and regulation of adult neurogenesis, prior work describing the vasculature structure within the DG has been largely qualitative. Still, these qualitative observations have revealed notable differences in vessel architecture by cell layer in the DG. The majority of cells in the DG are found in the densely packed granule cell layer (GCL). The less cell dense layers (ML and hilus (HL)) sit on opposing sides of the GCL, with the SGZ forming a thin cell layer between the GCL and hilus. In adult mice, the vasculature within the ML appears highly tortuous with many branching points (Licht and Keshet, 2015). The GCL layer contains visibly fewer vessel structures, and the vessels present are qualitatively linear (Lindhorst et al., 2012; Licht and Keshet, 2015). The vessels of the GCL then turn and run perpendicular in the SGZ. The vessels within the SGZ are densely packed and also often described as highly linear (Palmer et al., 2000; Lindhorst et al., 2012; Devasthali et al., 2025). HL vessels return to a more tortuous and dispersed path. While measures of vessel density across DG layers have been quantified in some studies, most descriptions of vessel tortuosity, angle and linearity have been purely qualitative to date.

A limitation to generating quantifiable measures of the blood vessel networks of the DG in prior work has been reliance on traditional thin, immunolabeled slices and 2-dimensional images. Features like the tortuosity, linearity and vessel diameter are inherently best appreciated in 3 dimensions (3D) and are therefore difficult to measure in a flattened image. Creating quantitative measures of these aspects of vessel structure can both deepen our understanding of the unique properties of the neurogenic niche and also lay the groundwork for future studies seeking to measure changes in vascular properties. To generate quantifiable measures of DG vasculature in 3D, we paired a post-mortem vessel casting technique with subsequent confocal imaging of cleared, thick coronal brain sections from adult mice. Images of large blood vessel networks were readily visible and quantified using 3D rendering software. Our approach replicated and quantified several previously well-established features of the DG vasculature, including several features that were previously only described qualitatively. Our data also revealed new unique and layer-dependent features of the neurogenic vascular niche (e.g. vessel diameter, branching) which have not been previously reported. Combined, our findings provide a quantitative basis for studying the vascular niche and expand the list of measurable features for future study. We also present details on our methodology, which uses widely-available tools, to facilitate use by others.

## Results

### Combination of Trypan blue and EZ clear facilitates blood vessel visualization for 3D rendering via IMARIS software

To identify all blood vessels in the dorsal DG, 9 week old male and female C57Bl/6J mice were perfused with a mixture of trypan blue and gelatin after a standard transcardial perfusion of phosphate buffered saline and paraformaldehyde (FIG 1. A-D). Trypan blue is an organic synthetic dye, which, due to its negative charge, does not pass through intact cell membranes. This property of exclusion from intact cell membranes has led to widespread use for distinguishing live versus dead (membrane-compromised) cells (Strober, 2015). Trypan blue has also been used to study the vasculature of the brain because when injected systemically, its cell-impermeant property coupled with the tight blood brain barrier results in its retention in blood vessels and exclusion from the brain interstitial tissue (Bentivoglio and Kristensson, 2014) (Fig 1B). In addition, while trypan blue is frequently visualized as a blue color in brightfield light, it also has reported fluorescence in the red to far red wavelength range (Avelar-Freitas et al., 2014; Kerschbaum et al., 2021) (wavelengths: 630 -720nm, expected peak: 660nm). We sought to leverage these properties of trypan blue (retention in the blood vessels and far-red fluorescence) for visualization and 3D reconstruction of intact vasculature. To further aid in retention of dye in the vessels, we mixed trypan blue with 1% bovine serum albumin (BSA). Trypan blue complexes with BSA, creating a much larger molecule which aids in the dye mixture’s retention within the blood vessels (Lang and Lasser, 1967). This dye-albumin mix was dissolved in 2% gelatin, which was transcardially perfused while warm to fill vessels, and then the whole mouse was rapidly cooled to allow the gelatin to solidify and preserve 3D structure of the blood vessels during post-processing. Fixed brains were then sectioned in thick, 1 mm coronal slices and cleared with the tissue clearing protocol, EZ Clear (Hsu et al., 2022)(Fig. 1C). A nuclear stain, Hoechst, was also added to visualize cell bodies within the DG, allowing segmenting of the DG into its layers based on cellular density (FIG 1D).

**Fig 1.**
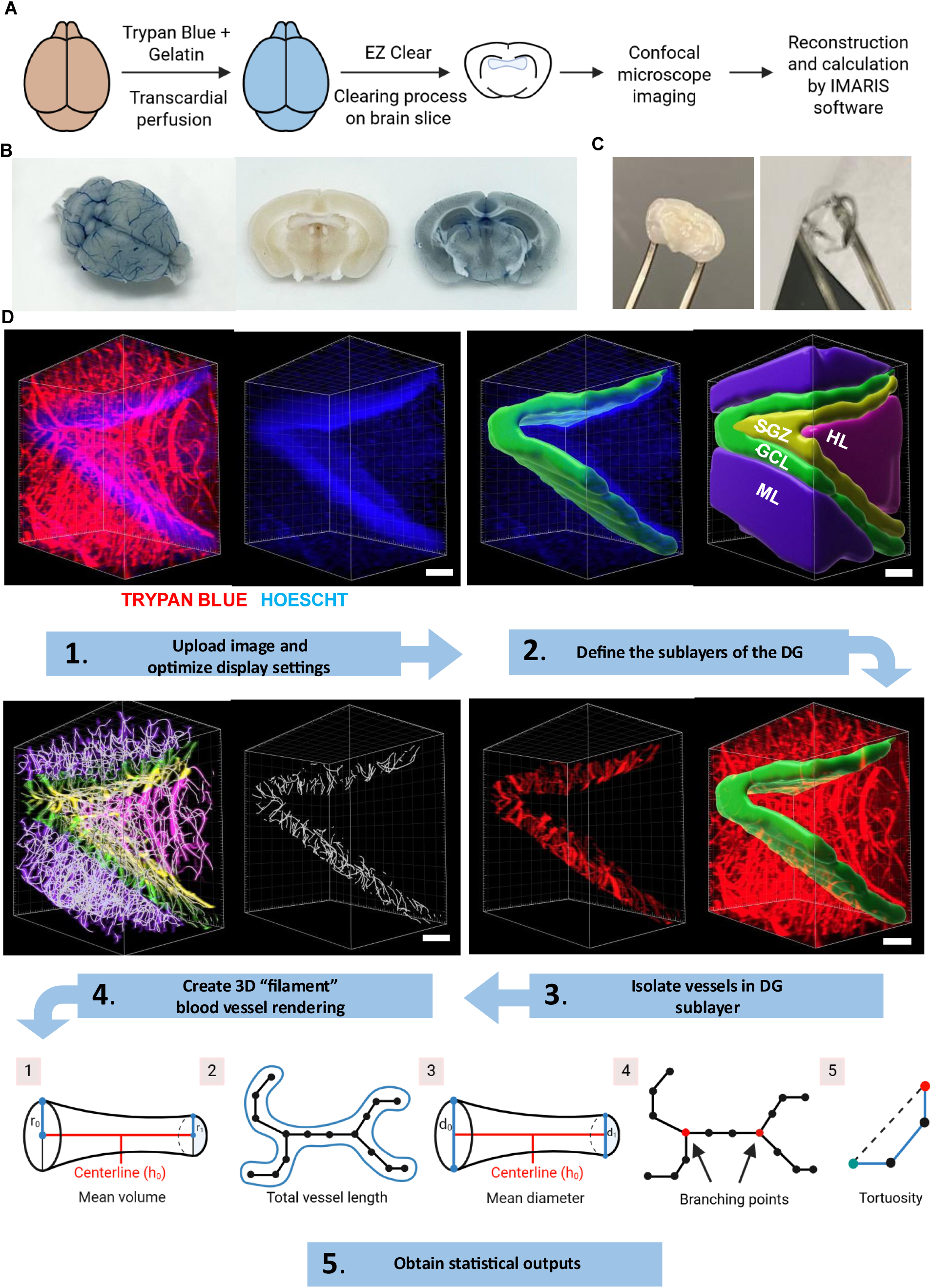
Blood vessels in the DG identified via trypan blue and EZ clear for 3D rendering in IMARIS software. (A) Experimental outline to characterize the DG blood vessels in 1 mm brain slices. (B) Image of whole mouse brain that has been perfused with paraformaldehyde (PFA), phosphate buffered saline (PBS) and the trypan blue/gelatin/BSA mixture beside an image comparing two 1 mm coronal brain slices. Within the slice comparison image; on the left is a 1 mm brain slice that has undergone a PFA + PBS perfusion and on the right is a 1 mm brain slice that has undergone a PFA + PBS + trypan blue/gelatin/BSA mixture perfusion. (C) Image of two coronal brain slices 1 mm thick. The slice on the left has not undergone the EZ clear process and the slice on the right has been cleared with the EZ clear tissue clearing process. (D) workflow of 3D vessel renderings and statistical output representations. Step 1: Unaltered z-stack of far-red fluorescence signal revealing trypan blue-filled vessels and UV-range signal showing Hoechst labeling in the DG. The Hoechst signal alone is shown on the right. Step 2: Hoechst label was used to define DG layers based on cell density and render them into surfaces within IMARIS. Left image shows an example of GCL surface pseudo-colored overtop of Hoechst labeling. Right image shows 3D rendered surfaces of the four layers of the DG. Step 3: DG layer surfaces were used alongside background subtraction to isolate the blood vessels within each DG layer. Right image shows an example of GCL surface pseudo-colored overtop of far-red fluorescence signal. Left image shows an example of the far-red fluorescence signal isolated within the GCL. Step 4: 3D rendered vessel filaments were created from trypan blue filled blood vessels. Left image shows an example of the blood vessel filaments pseudo-colored by DG layer and the GCL filaments are shown alone on the right in grey scale. Step 5: Statistical data on DG blood vessels was obtained through IMARIS. Numbered 1-5 are cartoons of the specified statistical outputs. From left to right: 1 is the mean volume, 2 is the total vessel length, 3 is the mean diameter, 4 is the branching points, 5 is the tortuosity. Scale = 100µm.

The DG of cleared slices was imaged in ultraviolet and far red wavelengths using confocal microscopy in z-stacks 269 to 447 µm thick, with a 1.5 µm optical slice interval (Fig 1A). Z-stacks were imported into IMARIS, a 3D imaging software. Figure 1D shows an example z-stack reconstructed in IMARIS. As step 1 of Fig 1D shows, distinct blood vessel structures were readily identifiable based on the endogenous far-red fluorescence of the trypan blue dye and the Hoechst counter-labeling revealed the densely cell-packed granular cell layer (step 1). To quantify vascular structure in the different layers of the DG, Hoechst labeling was used to guide creation of separate surfaces for the ML, GCL, SGZ and HL (Fig 1D, step 2). The far-red signal was then segmented by layer/surface using IMARIS’ “masking” tools (Fig 1D, step 3). IMARIS’ machine learning filament creator was then used to render the far-red signal of the trypan blue-filled vessels in each DG layer into filaments (Fig 1D, step 4). Along with the volume of the original segmented DG layer surfaces, the filament tool was used to quantify average volume, length, diameter, tortuosity and average number of branching points for the blood vessels within the DG (Fig 1D, step 4).

### Density of vessels varies by layer of the DG

Prior studies characterizing the vascular pattern of the DG layers have consistently found that the ML, HL, and SGZ have higher blood vessel density than the GCL (Palmer et al., 2000; Lindhorst et al., 2012; Dause et al., 2024; Devasthali et al., 2025). To determine whether these findings could be replicated using the EZ clear and subsequent 3D rendering method, we investigated the differences in density of blood vessels among the various DG layers. As expected, we found that the GCL had significantly lower vessel density than all other DG layers (all data reported as mean ± SEM; ML: 1.206 ± 0.1351 %; GCL: 0.4654 ± 0.06849 %; SGZ: 0.9481 ± 0.1299 %; HL: 0.9300 ± 0.1105 %) FIG 2 (A-D). In addition to volumetric measurements, we analyzed vessel length density (total vessel length over total layer volume) across the DG layers to better characterize vascular density patterns. Similar to volume-based measurements, vessel length density was significantly lower in the GCL compared to the ML (FIG 2D) (ML: 0.0006279 ± 7.225e-005 µm/µm^3^; GCL: 0.0003293 ± 5.442e-005 µm/µm^3^; SGZ: 0.0005419 ± 8.380e-005 µm/µm^3^; HL: 0.0005487 ± 6.418e-005 µm/µm^3^). Although it did not reach statistical significance after posthoc error correction, the SGZ and HL were also higher in vessel length density than the GCL, similar to the ML. Overall, these findings together suggest that our blood vessel visualization tools reliably capture the DG vascular architecture and provide quantitative validation of established anatomical observations of the DG vasculature.

**Fig 2.**
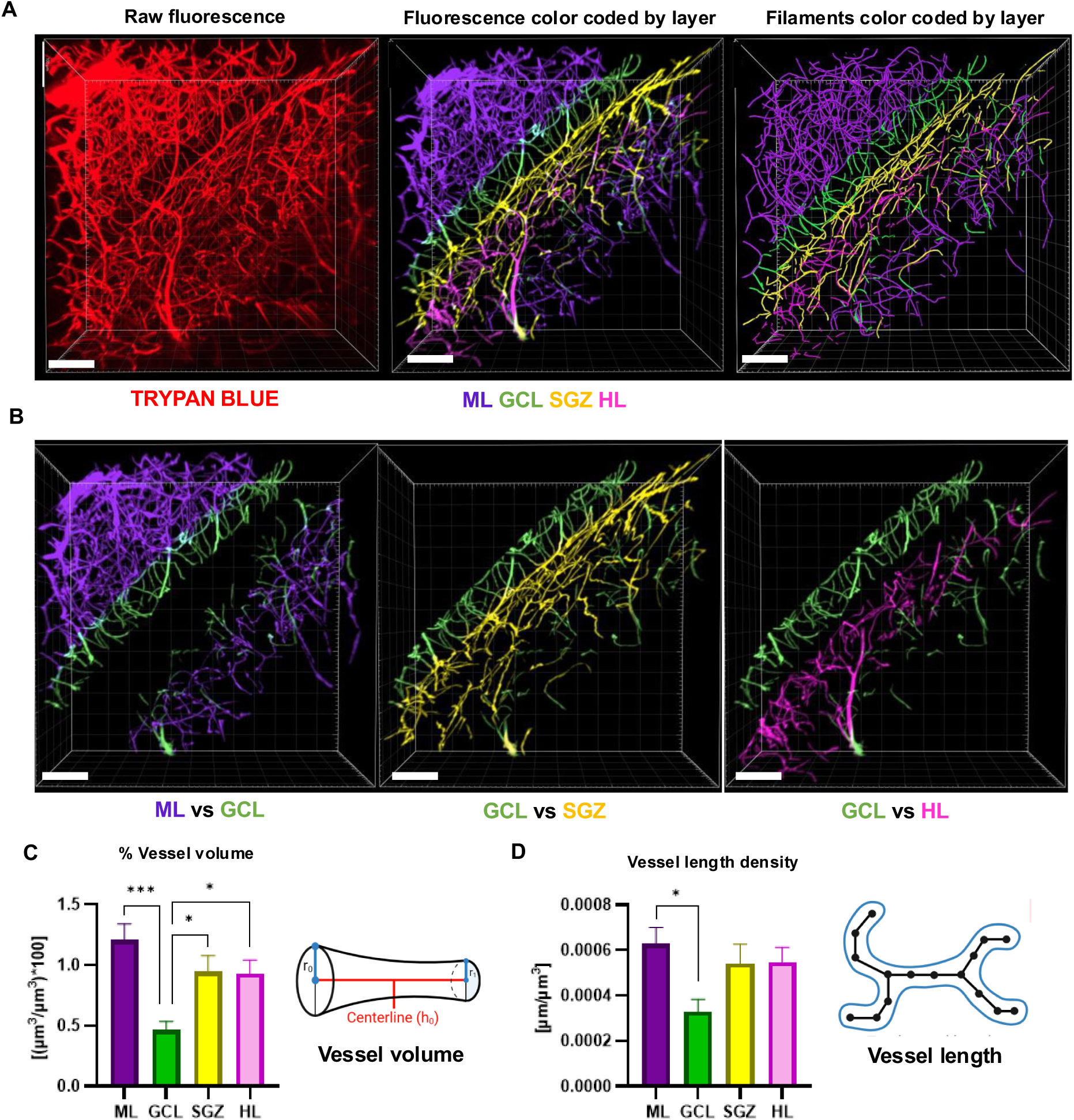
Percent volume density and length density of blood vessels is lower in the GCLvof the DG than other layers. (A) Representative far-red fluorescence images of DG blood vessels filled with trypan blue (left image), pseudo-colored vessels by DG layer (middle image), and pseudo-colored 3D filament renderings of the DG blood vessels (right image). Scale = 100 µm. (B) Representative image of pseudo-colored DG vessels comparing GCL with each other layer independently. ML and GCL vessels (left image), GCL and SGZ vessels (middle image), HL and GCL vessels (right image). Scale = 100 µm. (C) Mean ± SEM of percent vessel volume by DG layer (total volume of vessels/volume of layer)*100). One-way ANOVA p=0.0012, F _(3,24)_ = 7.301. ***p=0.0006 (GCL vs ML), *p=0.03 (GCL vs SGZ), *p=0.0384 (GCL vs HL) from Tukey’s multiple comparisons across layers. (D) Mean ± SEM of vessel length density by DG layer (total length of vessels/volume of layer). One-way ANOVA p=0.0345, F_(3,24)_ = 3.385. *p=0.0271 from Tukey’s multiple comparisons across layers. N = 7 mice

### Vessel branching is highest in the molecular layer of the DG

We next turned to quantifying aspects of vascular structure that have been previously less extensively studied in the adult DG niche and would benefit substantially from 3-dimensional visualization. First, we used IMARIS’ filament tool to quantify branching patterns of the vessels in each DG layer, a measure which to our knowledge has not been previously reported. The branching index was defined as the total number of branch points (BP) over the total length of the vessels in a particular DG layer. As expected from qualitative observations, the ML had a significantly higher branching index than any of the other DG layers (ML: 0.008756 ± 0.0005673 BP/µm; GCL: 0.003767 ± 0.0004989 BP/µm; SGZ: 0.005621 ± 0.0006669 BP/µm; HL: 0.005541 ± 0.0007009 BP/µm) (FIG 3 A-C). Although it did not reach statistical significance with post-hoc error correction, the GCL vessels exhibited less branch points per mm compared to the SGZ and HL, which is consistent with the qualitative differences that can be observed in the vascular architecture across these layers (FIG 3 A-C).

**Fig 3.**
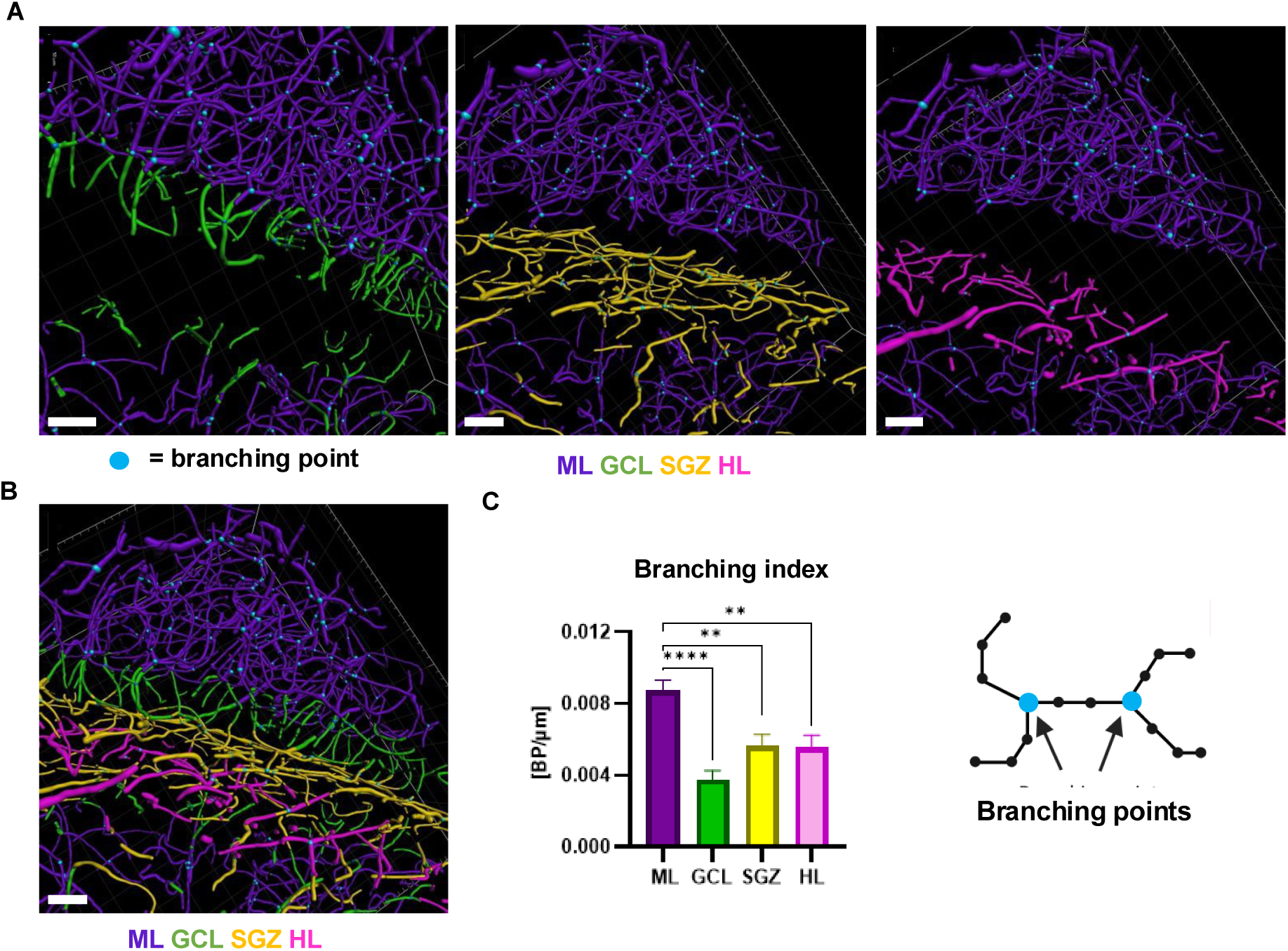
ML vessels have a higher branching index than GCL, SGZ, and HL. (A) Representative image of pseudo-colored 3D filament vessel renderings comparing ML and each other layer individually. Colored dots/spheres indicated below the first image represent branch points on a vessel. Shown are ML and GCL vessel renderings (left image), ML and SGZ vessel renderings (middle image), and ML and HL vessel renderings (right image). Scale is 50 µm. (B) Representative image of pseudo-colored vessel renderings for all four DG layers with indicated branch points. Scale is 50 µm. (C) Mean ± SEM of branching index by DG layer (number of branch points in layer/layer’s total vessel length). One-way ANOVA p<0.0001, F _(3,24)_ = 11.42. ****p<0.0001, **p=0.0071 (ML vs SGZ), **p=0.0057 (ML vs HL) from Tukey’s multiple comparisons across layers. N = 7 mice

### Vessels have a smaller average diameter in the granule cell layer of the DG than other layers

To our knowledge, vessel diameter also has not been previously quantified in the DG niche. Quantification of average vessel diameter using the IMARIS filament tool revealed that the GCL vessels had significantly smaller diameters on average as compared to the ML and SGZ (FIG 4 A-C) (ML: 4.798 ± 0.1481 µm; GCL: 4.150 ± 0.08593 µm; SGZ: 4.549 ± 0.09645 µm; HL: 4.466 ± 0.04486 µm). Hilar vessel diameter appeared to be intermediate between that of the GCL and SGZ (FIG 4 B).

**Fig 4.**
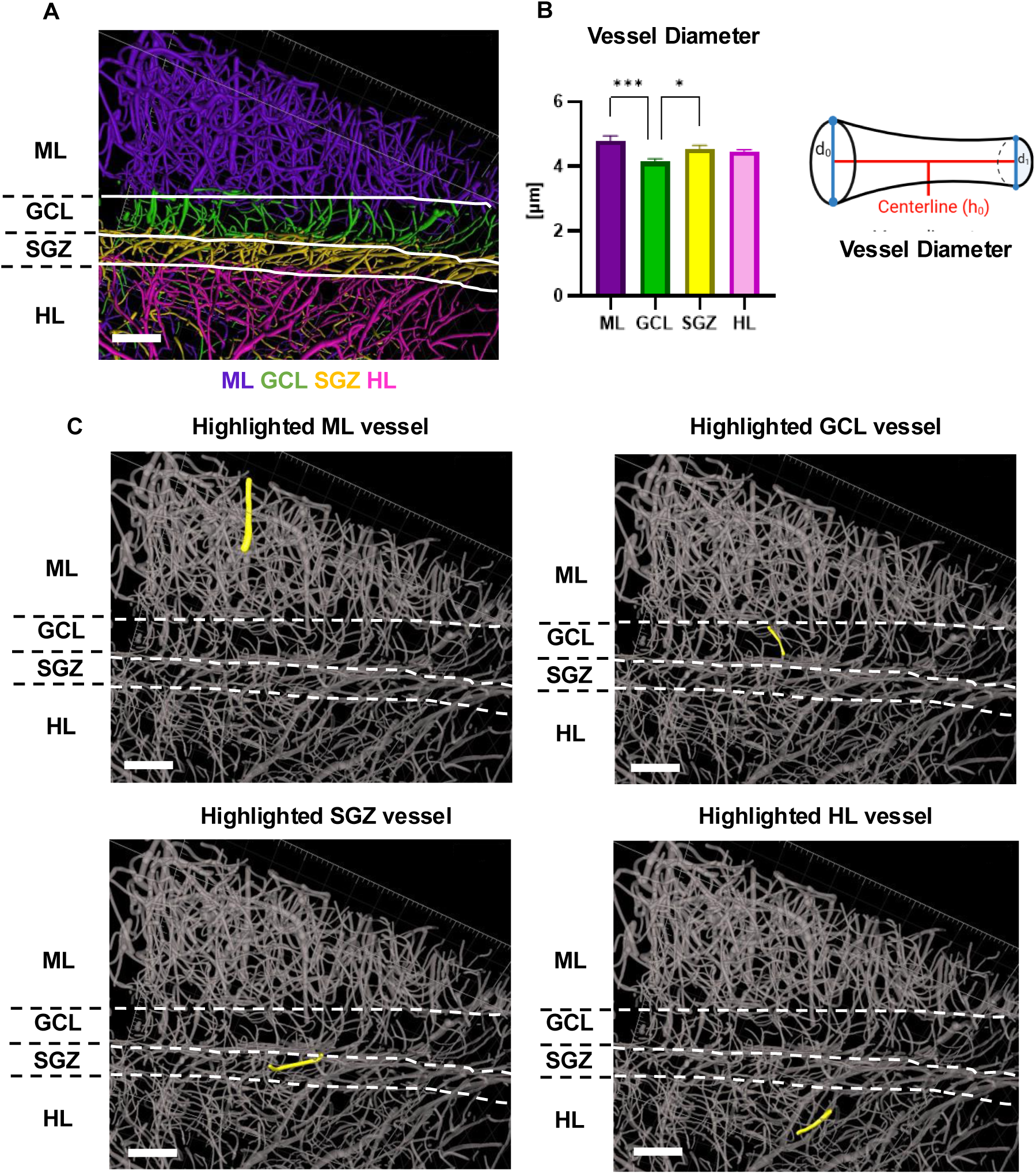
GCL vessels have a smaller diameter compared to ML and SGZ. (A) Representative image of pseudo-colored 3D filament vessel renderings. White lines across the image are separating the DG layers indicated by the label to the left of the image. Scale = 80 µm. (B) Mean ± SEM vessel diameter by DG layer. One-way ANOVA p=0.0015, F _(3,24)_ = 7.030. ***p=0.0007, *p=0.0456 from Tukey’s multiple comparisons across layers. N = 7 mice (C) Representative images of vessel renderings on a grey-scale. Dashed lines across the image are separating the DG layers indicated by the label to the left of the image. A 3D rendered vessel is highlighted in yellow in the ML (top left image), in the GCL (top right image), in the SGZ (bottom left image), and in the HL (bottom right image) to compare average vessel diameter. Scale = 80 µm.

### Molecular layer vasculature is more tortuous than other DG layers

The blood vessels of the SGZ have been widely cited to be exceptionally linear (Lindhorst et al., 2012; Licht and Keshet, 2015; Kim et al., 2021; Devasthali et al., 2025). This claim was originally based on qualitative observation of vessel structure, which reveals collections of vessels that appear to align to run parallel along the SGZ. Our previous study (Devasthali et al., 2025) quantified the planar orientation of the SGZ vessels and the perpendicular orientation of the GCL vessels. However, the angle of orientation and the linearity/tortuosity are distinct measurements. A vessel could be planar in orientation yet still follow a tortuous path. We therefore next sought to generate a quantitative measure of linearity/tortuosity across DG layers. Using the filament tool, linearity was measured on a scale of 0 (a circle) to 1 (a straight line). As expected from qualitative observations, the ML vessels were the most tortuous/least linear of all the layers, falling closer on average to 0 on the linearity index than other layers. With posthoc error correction, ML vessel linearity was significantly lower than the HL (Fig 5 A-C) (ML: 0.8604 ± 0.009121 (0-1 scale); GCL: 0.8951 ± 0.01293 (0-1 scale) SGZ: 0.8949 ± 0.008961 (0-1 scale); HL: 0.9000 ± 0.008024 (0-1 scale)). The other layers reflected the same overall pattern of having on average more linear vessels than the ML, though they were not significantly different compared to the ML after posthoc error correction. Intriguingly, GCL, SGZ and HL all showed largely similar linearity on average, suggesting that SGZ planarity is not especially unique among DG layers.

**Fig 5.**
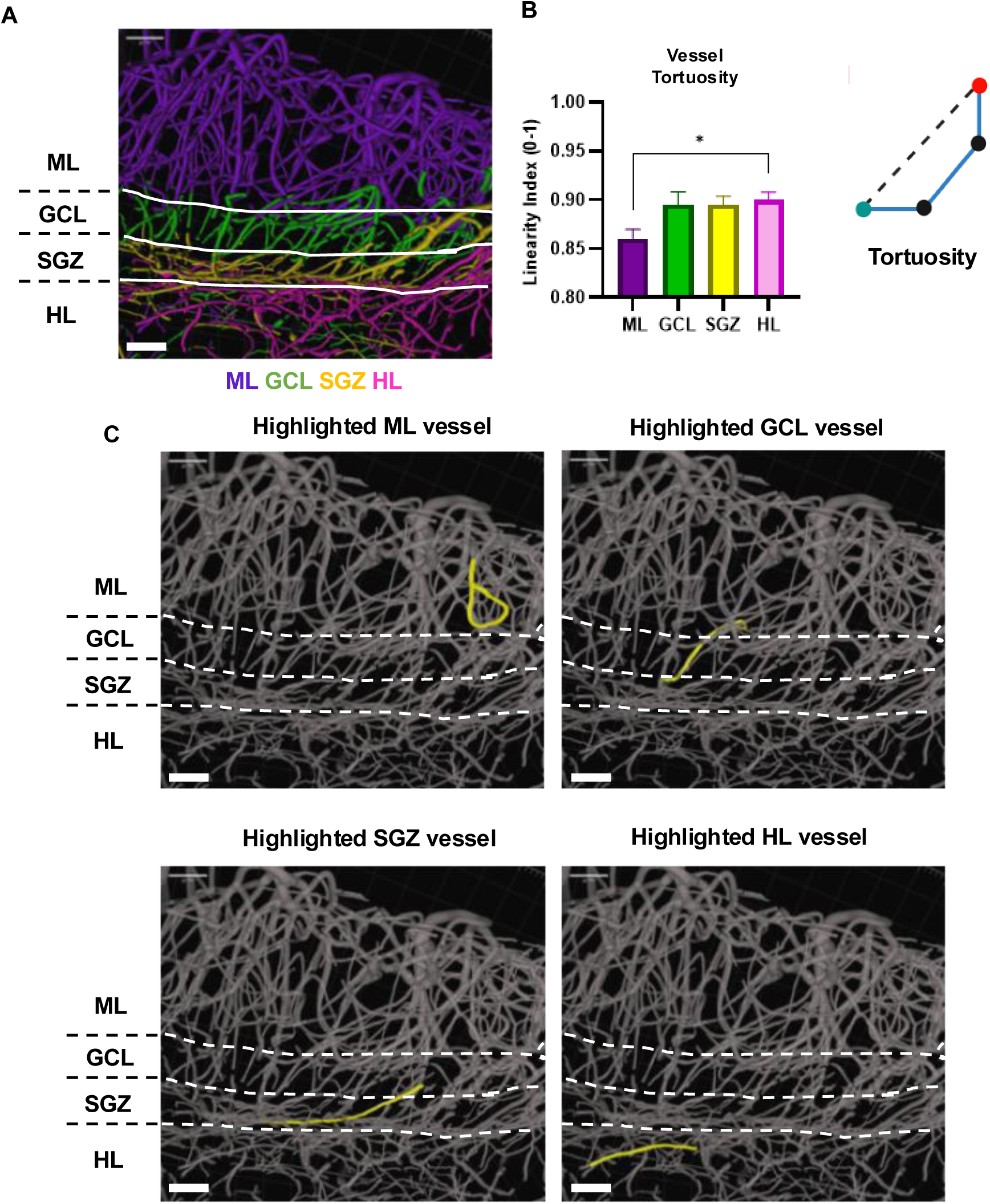
Vessels in the ML are highly tortuous. (A) Representative image of pseudo-colored 3D filament vessel renderings. White lines across the image are separating the DG layers indicated by the label to the left of the image. Scale = 50 µm. (B) Mean ± SEM linearity index by DG layer (vessels quantified on a scale of 0 (a circle) to 1 (a straight line)). One-way ANOVA p=0.0346, F _(3,24)_ = 3.381. *p=0.0442 from Tukey’s multiple comparisons across layers. N = 7 mice (C) Representative images of vessel renderings on a grey-scale. Dashed lines across the image are separating the DG layers indicated by the label to the left of the image. A 3D rendered vessel is highlighted in yellow in the ML (top left image), in the GCL (top right image), in the SGZ (bottom left image), and in the HL (bottom right image) to compare linearity. Scale = 50 µm.

## Discussion

Here, we provide a global characterization of the DG vascular niche in adult mice including features that had previously only been described qualitatively, as well as newly reported vessel features. Together, our findings both replicate previously established 2D tissue analysis of DG vessels and elucidate new distinctive layer-dependent DG vascular features.

Most prior work focused on the neurogenic vascular niche has used antibodies or lectin to label the endothelial cells that make up the vessel walls in thin (<60 µm thick) tissue slices then visualized that labeling in projected, 2D images (Palmer et al., 2000; Van Praag et al., 2005; Lindhorst et al., 2012; Dause et al., 2024, 2025; Devasthali et al., 2025). A smaller number of studies have used dyes or inks such as Evans blue, India Ink or trypan blue to fill the vessels, an approach more similar to the vessel casting technique we used here (Shimada et al., 1992; Jiang et al., 2014; Wälchli et al., 2015), but still typically visualizing the labeling in 2D images from thin slices. These prior approaches facilitated many qualitative assessments of the vascular niche, revealing several, key unique aspects of DG layer vessel structure. They also generated several useful vessel metric estimates, such as vasculature density and basic vessel branching/branching angle (Lindhorst et al., 2012; Dause et al., 2024; Devasthali et al., 2025). Because of the inherent limitations of visualizing 3D vessels in 2D projections, prior work has not generated measures of network-level features such as branching topology, true diameter, and path tortuosity. Here, we combined a simple dye+gelatin mixture for casting with tissue clearing of thicker slices (1 mm) and imaging on a standard confocal microscope to capture 3D vascular structure in the DG niche. This approach allowed us to reconstruct larger vessel networks in 3D using materials and tools that are readily available in many labs or core facilities. We were able to replicate key features of the neurogenic niche quantitatively, as well as quantify several new unique features of the DG layer vascular architecture. We suggest that these methods can be used to expand and refine the collection of features that can be visualized and quantified in future studies of the neurogenic vascular niche.

The SGZ vasculature has been described as particularly dense relative to neighboring DG layers (Palmer et al., 2000; Licht and Keshet, 2015), while the GCL has been reported to have sparser vessel density in comparison to the other DG layers (Lindhorst et al., 2012; Licht and Keshet, 2015). Quantifications using density of area of immunolabeling for vessel protein markers in traditional 2D images have confirmed the qualitatively low vessel density of the GCL, while the relative differences between other layers were less clear and consistent (Lindhorst et al., 2012; Dause et al., 2024; Devasthali et al., 2025). Similar to prior work in traditional 2D microscopy, we found that the GCL exhibited significantly lower vessel density than all other DG layers when rendered in 3D from fluorophore-filled vessels in thick, cleared sections. As noted in prior work, it seems likely that the high density of granule cell neurons in the GCL leaves less physical space available for blood vessels, compared to the more cell-sparse ML and HL. Although the SGZ has previously been described as the most vascularly dense region of the DG (Palmer et al., 2000; Licht and Keshet, 2015)), it is notable that prior work using quantitative measures of vessel marker immunolabeling in 2D microscopy revealed, at best, subtle differences in vessel density between the SGZ and HL/ML. We similarly found that SGZ density was elevated only relative to the GCL and was not significantly higher than that of the other DG layers. The qualitative description of the SGZ as having uniquely high vessel density may reflect a visual effect driven by the uniform, planar organization of SGZ vessels, in contrast to the more tortuous and randomly oriented vessels of the other DG layers. These findings underscore the value of quantitative measures over qualitative ones, which can be misleading at times.

The differing levels of linearity and tortuosity across DG layers is one of the more visually striking features of the DG vasculature, although this aspect of the vascular niche has thus far only been characterized qualitatively. Prior descriptions indicate that DG capillaries originate from arterioles at the hippocampal fissure, descend perpendicularly through the GCL, and then curve to run parallel to the SGZ in a distinctly linear fashion (Lindhorst et al., 2012; Licht and Keshet, 2015; Kim et al., 2021; Devasthali et al., 2025), while ML and HL vessels, though less well described in the literature, follow a more tortuous and winding path. Our visualization approach enabled quantification of this feature by defining a linearity index ranging from 0 (a circle) to 1 (a straight line). On average, ML vessels were substantially more tortuous, falling closer to 0 on the linearity index than vessels in the other DG layers, consistent with prior qualitative descriptions of the SGZ and GCL as particularly linear relative to other layers. Somewhat unexpectedly, HL vessels were also relatively linear, despite qualitative descriptions characterizing them as non-linear. This discrepancy may stem from the random directionality of HL vessels, in contrast to the single uniform orientation of GCL and SGZ vessels within their respective layers. Again, these results underscore the value of quantification over qualitative assessments of vascular structure.

The functional significance of linear versus tortuous vasculature across DG layers remains unclear. One possible explanation for the pronounced tortuosity of ML vessels is the cellular composition of the region. The ML is composed primarily of dentate granule neuron dendrites rather than densely packed cell bodies. More physical space is therefore available for vessels to wind through the extracellular space than in more cell-dense layers such as the GCL or SGZ. Alternatively, ML vessel tortuosity may reflect the surface-contact demands of dendritic trees that fill the middle and outer ML and NSC bushy terminals, which contact vessels in the inner ML (Moss et al., 2016; Licht et al., 2020). A more winding vessel path could provide more opportunities for proximity to these cellular processes. The linearity and uniformity of SGZ vessels, meanwhile, has been hypothesized to facilitate the tangential migration of neuroblasts (Sun et al., 2015), providing a clear migration path parallel to the GCL for differentiating cells to travel before dissociating to migrate into the deeper GCL layers where they will adopt more mature neuronal features. The functional roles of tortuous versus linear vasculature in the DG remain a topic for future study.

The average diameter of DG vessels by layer has not previously been reported qualitatively or quantitatively. Our findings show that GCL vessels had, on average, a smaller diameter than vessels in the other DG layers, which may again be attributable to the dense cellular composition of the GCL. The tightly packed region may impose spatial constraints that demand smaller vessel diameters. Our visualization method was particularly critical for generating accurate measurement of layer-dependent vessel diameter throughout the DG. Vessel-casting with trypan blue dye, albumin, and gelatin rapidly filled the vessels and produced a solid, core-like structure. This approach stands in contrast to the hollow lumen typically visualized via endothelial immunostaining, which can be more subject to distortion (stretch, compression) during tissue processing. Additionally, the thick-tissue nature of our imaging method allowed more vessels to be analyzed in their entirety. The vessel cutting common to traditional thin-tissue analysis can make it difficult to capture enough intact vessel length to measure full diameter.

Layer-dependent vessel branching in the DG has also been comparatively understudied in prior work. A small number of studies have reported a pattern of low branch-point density in GCL vessels alongside similarly high branching in SGZ and ML vessels (Lindhorst et al., 2012; Devasthali et al., 2025), though branching was not directly compared across layers in the adult DG. In alignment with this prior work, we found that the ML exhibited a significantly higher branching index than all other DG layers. SGZ branching, however, was lower than ML and more similar to GCL and HL. This novel result may reflect differences between traditional thin-tissue 2D analysis and our thick-tissue 3D approach. Whereas 2D data show the SGZ and ML as similar in branching index, 3D imaging reveals a clear and significant elevation in ML branching. This discrepancy may arise because the linear, planar organization of SGZ vessels makes branch points easier to capture within a single thin tissue slice, whereas the greater tortuosity of ML vessels means they are more likely to weave in and out of a thin slice, causing individual bifurcated vessels to be miscounted as separate, unbranched vessels.

There are various approaches to imaging brain vasculature in three dimensions that have been used in prior studies of a variety of brain regions. Studies vary in several major elements: 1) how the vessels are labeled, 2) what signal is used to visualize vessels and 3) what tool is used to capture images of large volumes of tissue.

Vessel labeling for 3D visualization is most commonly achieved through one of two primary methods: filling the lumen of the vessels via vessel casting, or labeling the endothelial cells on the walls of the vessels via immunolabeling or lectin labeling (Kirst et al., 2020; Todorov et al., 2020). We chose to use vessel casting here because it allowed us to visualize the continuous interior volume of the blood vessels rather than relying on endothelial cell labeling, which can be inconsistent and leaves the vessel interior hollow. In our previous attempts, a hollow vessel interior made 3D rendering and subsequent analysis in IMARIS difficult. Whether casting or immunolabeling is used, visualization of large, 3D structures typically requires some kind of tissue clearing. We chose the EZ Clear protocol, which uses tetrahydrofuran to delipidate the tissue and render it transparent by matching the refractive index of the tissue to that of the imaging media (Hsu et al., 2022). It is relatively inexpensive, preserves fluorophores, and does not require specialized equipment. Many alternative approaches are available to render thick brain tissue transparent, including iDISCO (Renier et al., 2014), CLARITY (Chung et al., 2013), and CUBIC (Susaki et al., 2014). Each clearing method carries its own tradeoffs. iDISCO uses hazardous reagents and can deform tissue or quench fluorophores (Ueda et al., 2020). CLARITY, though less hazardous, requires specialized equipment (Chung et al., 2013; Tomer et al., 2014). Finally, CUBIC is most similar to the EZ Clear method, as it avoids both hazardous materials and equipment costs, but it requires long clearing times (Susaki et al., 2014, 2015). EZ Clear therefore offered the optimal route for clearing tissue while preserving fluorophores contained in cast vessels.

Vessel visualization, whether generated via casting or immunolabeling, is most commonly done via fluorescence microscopy. To make our vessel casting fluorescent, we used the fluorescent signal from trypan blue dye, which was mixed with BSA and gelatin to help solidify the dye in the vessels. Trypan blue is a relatively inexpensive and commonly available dye, which made it practical choice to use as opposed to a more expensive fluorophore-conjugated albumin like FITC-albumin. Some non-fluorescent alternatives for vessel casting signal media include resin or iodine. These casting media are optimized for X-ray imaging and while this approach can generate images at greater depth, they require micro-CT hardware, and resin casting can be technically difficult, particularly when attempting to co-label with a fluorescent marker such as Hoechst (Starosolski et al., 2015; Quintana et al., 2019).

Finally, there are 2 primary tools available for capturing 3D images of cleared tissues: confocal microscopes and lightsheet microscopes. Lightsheet microscopes can rapidly image large samples, including entire brains, but are highly specialized and not available to many labs. In addition, resolution on many lightsheet microscopes, particularly at the low magnifications used to image large samples, can be insufficient to see individual microvessels (Santi, 2011; Ueda et al., 2020). Confocal microscopes offer much greater spatial resolution than lightsheet microscopes and are more widely available across labs. They are, however, limited in the thickness of samples that can be imaged effectively. To capture microvessel structure, we chose to image a singular thick brain slice, approximately 1 mm, using a confocal microscope. While this approach prevents imaging of the entire DG in one sample, it offers a practical middle-ground where a large sampling of vessel network structure is captured in most of the dorsal section of the DG.

There are several limitations to the current study. First, while the trypan blue dye mixture allowed for visualization of a continuous vessel structure, it only provides an estimate of interior vessel diameter. The endothelial cells lining the outer vessel wall remain unobserved in our approach. Incorporating endothelial cell thickness into vessel diameter measurements may provide a more complete picture of how other cell types, like NSCs and neuroblasts, interact with the vasculature, particularly in the context of RGL-NSCs wrapping around iML vessels. A future study combining the advantages of vessel casting with endothelial cell staining would help address this limitation. Second, vessel casting relies on effective perfusion and dispersion of the dye/gelatin mixture through vessels. Errors during perfusion or collapse of blood vessels could leave some blood vessels unfilled and therefore unavailable to measure during analysis. A future study with advanced imaging and combination of vessel casting and endothelial cell staining could also solve this issue by checking for vessels that were positive for endothelial cell stains and negative for trypan blue in the lumen. Finally, our study focused solely on the visualization of blood vessels but did not show how other cells associate with those blood vessels. Additional visualization of NSCs alongside the vasculature in 3D would provide insight into how vessels interact with the neural stem cell niche more broadly, such as through vessel wrapping in the iML (Moss et al., 2016; Licht et al., 2020) or the positioning of SGZ cell bodies relative to vessels (Palmer et al., 2000; Dause et al., 2024; Devasthali et al., 2025). However, given the small scale of such interaction points, it seems likely that these cell-to-vessel associations are well captured using confocal imaging of thin sections immunolabeled to visualize NSC features. Thick sections and larger scale 3D vessel reconstructions may not be necessary to study such fine features of cell morphology.

In conclusion, our study provides an in-depth quantitative characterization of the DG vascular architecture and establishes a methodology that can be readily used for various unique vessel measurements in future studies. Understanding the anatomy of the DG vascular niche provides insight into how NSCs and neurogenesis persist in their endogenous hippocampal microenvironment. Studying these endogenous niches can reveal general mechanisms for supporting stem cell survival in adult brain tissue, a key step for developing regenerative medicine treatments for various neurological diseases and disorders.

## Methods

### Mice

All of the mice were in standard ventilated cages on a 12/12 hour light/dark cycle with ad libitum access to food and water. NestinGFP mice (#033927) were used from Jackson Labs and bred with C57Bl/6J in The Ohio State University Psychology Building mouse vivarium. Both NestinGFP-positive (hemizygous) and NestinGFP-negative littermates were used in this study. The pups were weaned at 3 weeks of age and housed in same-sex groups of 2-5. All animal use was in accordance with institutional guidelines approved by The Ohio State University Institutional Animal Care and Use Committee. Male and female mice were included and data from both sexes were combined for the main analyses (see Statistical analysis).

### Tissue collection / Trypan blue - BSA - gelatin dye mixture administration

9 mouse litters were randomly selected from 9 weeks (n = 7, 3M/4F) of age. For tissue harvesting mice were anesthetized using a mixture of 87.5 mg/kg ketamine and 12.5 mg/kg xylazine. Mice were then transcardially perfused with 0.1M PBS then followed by chilled 4% paraformaldehyde (Fisher #AC169650010) in 0.1 M phosphate buffer to partially fix the brain and remove blood. The mice were then transcardially perfused with a final mixture of 2% gelatin (Sigma-Aldrich #G1890) w/v, 1% BSA (Cell Signaling Technology #9998S) w/v, 50% Trypan blue (Fisher #15250-061) v/v, 50% 1X PBS v/v. The mice were then submerged in an ice bath for 5-10 minutes to allow the gelatin/trypan blue mixture to solidify in the blood vessels.

### Tissue processing / EZ clear protocol

Mouse brains were sliced immediately after perfusion using an ice-cooled brain matrix in a 1:1 series of coronal 1mm thick sections, measured using calipers. Sections were then post-fixed in 4% paraformaldehyde (Fisher #AC169650010) in 0.1M phosphate buffer at 4°C for 24 hours in the dark. The sections were then delipidated with 50% v/v Tetrahydrofuran (Sigma-Aldrich #186562) in 50% v/v deionized water and placed overnight at room temperature in the dark and in a fume hood. Sections were then placed in 0.05% w/v Sodium Azide (Sigma-Aldrich #S2002) in 1X PBS in the dark at 4°C to store until they were ready for imaging. The sections were then placed in 1hr rinses 2x in deionized water, followed by a third rinse with Hoechst 33342 (1:2000 in deionized water, Fisher #H3570). The EZ clear imaging media was then made with 7M urea (Fisher #42458), 0.025% sodium azide, 80% Nycodenz (Accurate Chemical and Scientific Corp. #AN1002423), in 2X PBS. Sections were then placed in the EZ clear imaging media for 24hrs at 4°C in the dark. A custom-built chamber was created to mount the sections in the EZ clear imaging media for imaging on the microscope. For long term storage the sections were left in the EZ clear imaging media in the dark at 4°C.

### Microscopy

The DG was imaged using the Zeiss LSM 900 Airyscan 2 at 20x magnification in 200-600 µm z-stacks with a 1.5 µm interval between optical sections. Two channels were utilized, Hoechst nuclear labeling (405 nm excitation) and trypan blue far-red fluorescence (640 nm excitation). Z stacks were converted to IMARIS files and processed in IMARIS as 3D renderings of the DG.

### Image quantification / IMARIS

Hoescht-labeled DG layers were manually contoured in Imaris (Imaris 10.2, Oxford Instruments) using the Surfaces module’s contour drawing tool. This step generated a layer-specific 3D surface for each region and was done across the z-stack at roughly 5–10 μm or 40–50 μm intervals depending on the layer thickness. The Trypan channel (640 nm excitation, ∼650–700 nm emission) was then masked to each layer’s surface to isolate further vessel reconstruction to that specific DG layer. To isolate vessel signal from background, a new surface was generated from the masked channel using background subtraction (largest vessel diameter set per image based on the maximum measured vessel width in that stack), and this surface was used to mask the channel a second time. The blood vessels were then rendered from the double-masked channel using the Filament tool’s autopath (loops, no soma, no spines) algorithm with multiscale seed point detection. Seed points and filament segments were created using the built-in machine-learning train-and-predict classifier to remove falsely detected points and non-vessel connections (“loops” from falsely detected vessel connection points), followed by a size threshold to exclude small filament fragments (the smallest filaments were artifacts from the masked channel). Filament statistics like length, diameter, linearity, branch points, etc. were collected per DG layer using the “Statistics” module and later used for analysis.

### Statistical analysis

Statistical analyses were performed using GraphPad Prism (version 10.0.3 (275), www.graphpad.com). Data for each vascular metric was assessed by ordinary one-way ANOVAs for each DG layer followed by Tukey’s multiple comparisons test. To assess potential sex differences, each measure was also analyzed by two-way repeated measures ANOVA (layer x sex). No main effects of sex or layer x sex interactions were observed for any measure except vessel diameter, which showed a modest layer x sex interaction (F(2.40, 12.02) = 3.97, p = 0.042). However, Šídák’s multiple comparisons revealed no significant sex differences within any DG layer (all p > 0.10), and the GCL had the smallest average vessel diameter in both sexes (Fig. S1). Data were therefore collapsed by sex for the main analyses. Data were approximately normally distributed. All statistical tests were two-tailed with α = 0.05.

### Reagents table

| Reagent or resource | Source | Identifier |
| --- | --- | --- |
| Antibodies/Dyes |  |  |
| Hoechst 33342 (1:2000) | Fisher | Cat#H3570 |
| Trypan blue | Fisher | Cat#15250-061 |
| Chemicals and reagents |  |  |
| Paraformaldehyde, 4% | Fisher | Cat#AC169650010 |
| Gelatin | Sigma-Aldrich | Cat#G1890 |
| BSA | Cell Signaling Technology | Cat#9998S |
| Tetrahydrofuran | Sigma-Aldrich | Cat#186562 |
| Sodium Azide | Sigma-Aldrich | Cat#S2002 |
| Urea | Fisher | Cat#42458 |
| Nycodenz | Accurate Chemical and Scientific Corp. | Cat#AN1002423 |
| Experimental models: Mouse models |  |  |
| NestinGFP | Jackson Laboratory | #033927 |
| Wild-type C57BL/6J | Jackson Laboratory | #000664 |
| Software and algorithms |  |  |
| Prism | GraphPad | www.graphpad.com |
| IMARIS | Oxford Instruments |  |

## Data availability

The datasets generated during and/or analyzed during the current study are available from the corresponding author on reasonable request.

## Acknowledgements

This work was funded by R01NS124775 from the National Institute of Neurological Disorders and Stroke to EDK. We thank the Neuroscience Imaging Core for access to imaging resources. Imaging was performed on a Zeiss LSM 900 Airyscan 2 confocal microscope, which was purchased with NIH Shared Instrumentation Grant S10 OD026842. Diagrams were created with BioRender.com.

## Author contributions

EDK and IC conceived and designed the study. EDK, IC, and BMS were involved in animal care, experimental procedures, and data collection. IC and BMS collected data and performed the data analysis. IC wrote the first draft of the manuscript and EDK edited and critically reviewed the manuscript. All authors reviewed the results and approved the final version of the manuscript.

## Additional information

The authors declare no competing interests.

