## Supplemental Figure 1 for "Classic and novel features of the dentate gyrus vascular niche quantified through 3D vessel reconstruction"

### Supplementary materials:

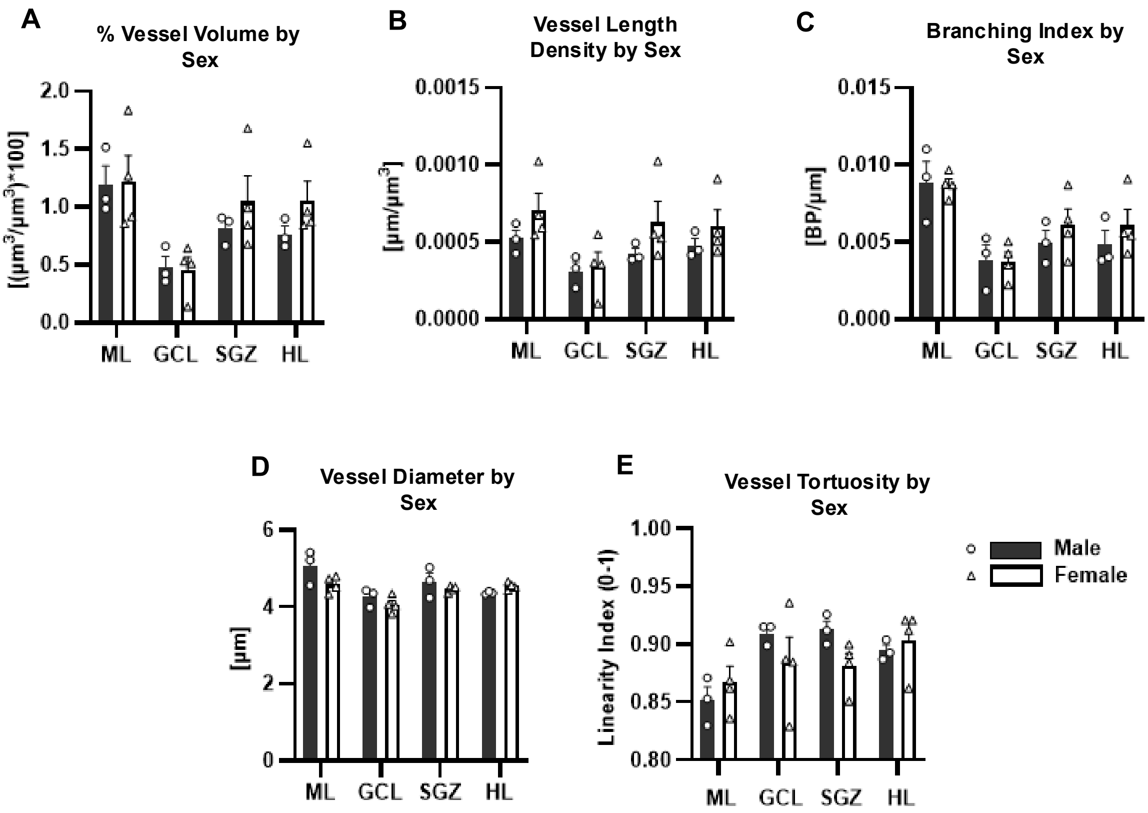


**Supplemental Fig S1. DG vascular measures by sex.** Mean ± SEM of (A) percent vessel volume density, (B) vessel length density, (C) vessel branching index, (D) average vessel diameter, and (E) vessel linearity index in the ML, GCL, SGZ, and HL of male (filled bar) and female (open bar) mice. Points represent individual mice (individual male mice represented by circle points, individual female mice represented by triangle points). Data were analyzed by two-way repeated measures ANOVA (layer × sex) with Geisser-Greenhouse correction. No significant main effects of sex were observed for any measure (A: $F(1,5) = 0.3778$, $p = 0.5657$; B: $F(1,5) = 1.092$, $p = 0.3439$ ; C: $F(1,5) = 0.2767$, $p = 0.6214$; D: $F(1,5) = 1.365$, $p = 0.2954$; E: $F(1,5) = 0.5244$, $p = 0.5014$). A significant layer x sex interaction was observed only for vessel diameter (D; $F(2.40, 12.02) = 3.97$, $p = 0.042$), but Šídák's multiple comparisons revealed no significant sex differences within any layer (ML $p = 0.60$, GCL $p = 0.80$, SGZ $p = 0.94$, HL $p = 0.11$). N = 7 mice (3 male, 4 female).
